# PARG inhibition induces replication catastrophe in ovarian cancer cells with down-regulated DNA replication genes

**DOI:** 10.1101/2020.07.13.199968

**Authors:** C. Coulson-Gilmer, R.D. Morgan, L. Nelson, B. Barnes, J.C. McGrail, S.S. Taylor

## Abstract

The discovery that PARP1/2 inhibitors selectively kill *BRCA* mutant cells has led to a paradigm shift in the treatment of women with homologous recombination (HR)-deficient high-grade serous ovarian cancer (HGSOC), driving unprecedented improvements in progression-free and, more recently, overall survival. However, because most HGSOC cases are not HR-defective, and are therefore unlikely to benefit from PARPi-based therapies, additional strategies will be required to improve outcomes for women with HR-proficient disease. To develop novel therapeutic strategies, considerable attention is now being focused on inhibitors targeting PARG, the poly(ADP ribose) glycohydrolase that counterbalances PARP1/2 activity. Here we characterise ten ovarian cancer cell lines in response to the PARG inhibitor PDD00017273, hereafter PARGi. We demonstrate that six lines are resistant while four are sensitive, and that sensitivity correlates with several markers of persistent DNA replication stress, DNA damage and replication catastrophe, namely the accumulation of asymmetric DNA replication fibres, γH2AX and RPA foci, KAP1 and Chk1 phosphorylation, a pre-mitotic cell cycle block and, following prolonged exposure, a pan-nuclear γH2AX phenotype that indicates RPA exhaustion. We demonstrate that PARGi-sensitive cell lines have down-regulated DNA replication genes, including components of the fork protection complex, namely *TIMELESS*, *TIPIN* and *CLASPIN*. These observations suggest that a subset of HGSOC may respond to PARG inhibitors and that “*replication stress*” gene expression signature could serve as a predictive biomarker to guide the design of clinical trials.

## Introduction

Ovarian cancer is a leading cause of female cancer-related death, accounting for ∼185,000 deaths globally in 2018 (Bray et al., 2018; Ferlay et al., 2018). The most prevalent subtype, high-grade serous ovarian carcinoma (HGSOC) is particularly lethal because it develops rapidly and often presents at an advanced stage (Lheureux et al., 2019). Treatment options are limited; typically, cytoreductive surgery and paclitaxel/platinum-based chemotherapy, maintenance therapy and hormone antagonists (Ledermann et al., 2013; Lheureux et al., 2019; NCCN Guidelines, 2020). While many patients initially respond well, most develop recurrent disease, yielding 10-year survival rates of only ∼35% (Bowtell et al., 2015).

HGSOC is characterised by ubiquitous *TP53* mutation and extensive copy number variation (CNV) (Ahmed et al., 2010; Ciriello et al., 2013). *BRCA1/BRCA2* are inactivated in ∼20% of cases, leading to HR deficiency (HRD) (Cancer Genome Atlas Research, 2011), however DNA damage repair (DDR) defects are more widespread (Patch et al., 2015; McCormick et al., 2017). Extensive CNV implies chromosome instability (Sansregret et al., 2018), and indeed, HGSOC is one of the most chromosomally unstable cancers (Bowtell et al., 2015; Nelson et al., 2020).

While precision medicine is revolutionising cancer treatment, this paradigm is challenging in HGSOC due to the paucity of actionable driver mutations. Other therapeutic strategies are therefore required and indeed, HRD has opened up an alternative, namely synthetic lethality. This approach was pioneered by the ability of PARP inhibitors (PARPi) to selectively kill *BRCA*-mutant cells (Bryant et al., 2005; Farmer et al., 2005), and these drugs are now yielding major benefits for women with *BRCA*-mutant ovarian cancer (Moore et al., 2018; Coleman et al., 2019; Gonzalez-Martin et al., 2019; Ray-Coquard et al., 2019).

While only ∼20% of HGSOC cases have *BRCA1/BRCA2* mutation, a further ∼30% are predicted to be HRD due to other oncogenic lesions and thus might also benefit from PARPi (Cancer Genome Atlas Research, 2011; Ray-Coquard et al., 2019). However, this still leaves up to ∼50% of cases that are HR-proficient and unlikely to benefit from a PARPi-based strategy. Thus, additional strategies will be required to improve outcomes for women with HR-proficient disease.

To develop novel therapeutic strategies targeting HR-proficient cancers, considerable attention is now being focused on inhibitors of PARG, the poly(ADP ribose) glycohydrolase that counterbalances PARP1/2 activity (Koh et al., 2004; Gravells et al., 2017; Chen and Yu, 2019; Gogola et al., 2019; Pillay et al., 2019). Thus far, we have characterised PDD00017273, hereafter PARGi, a quinazolinedione that inhibits PARG with an *in vitro* IC50 of 26 nM (James et al., 2016). Upon analysing a panel of six ovarian cancer cell lines in response to PARGi and the PARP1/2 inhibitor Olaparib, we discovered that while OVSAHO, COV318, COV362 and CAOV3 proliferated in both inhibitors, Kuramochi and OVCAR3 displayed differential sensitivities; Kuramochi proliferation was only suppressed by PARGi, while OVCAR3 proliferation was only suppressed by PARPi (Pillay et al., 2019).

Sensitivity of Kuramochi to PARGi was accompanied by pan-nuclear γH2AX staining, indicating replication catastrophe (Toledo et al., 2017), and indeed a synthetic lethal siRNA screen identified several DNA replication genes, including *TIMELESS*, that – when inhibited – sensitized OVCAR3 to PARGi. Analysis of DNA replication fibres showed that PARGi induced fork asymmetry in Kuramochi but not OVCAR3, leading to the conclusion that Kuramochi cells have an underlying DNA replication vulnerability that causes frequent fork stalling and a thus a dependency on PARG to re-start stalled forks by reversing PARP1-mediated inhibition of RECQ1 (Pillay et al., 2019).

The notion that DNA replication vulnerabilities might explain PARGi sensitivity led to the identification of additional sensitive cell lines. Interrogation of Cancer Cell Line Encyclopaedia (CCLE) data identified several ovarian cancer cell lines with down-regulated DNA replication genes, namely Kuramochi, HS571T, RMG1, OV56, OVMANA and OVISE (Pillay et al., 2019). While HS571T is no longer available for research purposes, we obtained RMG1, OV56, OVMANA and OVISE, and showed that RMG1 and OVMANA are PARGi sensitive. This suggests that a DNA “*replication stress*” gene expression signature might have potential as a predictive biomarker of PARGi sensitivity. Importantly, like Kuramochi, RMG1 showed differential sensitivity to PARGi and the PARPi olaparib, and while OVMANA were more sensitive to PARGi than PARPi, the distinction was less clear-cut.

These observations indicate that PARG inhibitors may open up new opportunities to treat HR-proficient ovarian cancers, which are unlikely to respond to PARP inhibitors; however, a number of questions remained unanswered. Firstly, is the PARGi sensitivity exhibited by different cell lines mediated via the same mechanism? While PARGi prevents proliferation of RMG1 and OVMANA, whether this was due to the replication catastrophe phenomenon exhibited by Kuramochi cells was not established. Secondly, does PARGi sensitivity correlate with down-regulated DNA replication genes? While the CCLE data indicated that DNA replication genes are down-regulated in OV56 and OVISE, their PARGi sensitivity was unclear (Pillay et al., 2019). If resistant, the utility of a “*replication stress*” gene expression signature as a predictive biomarker of PARGi sensitivity is uncertain.

To address these two questions, we set out to perform a detailed analysis of a panel of ovarian cancer cell lines to determine (a) whether PARGi sensitivity is via a common replication catastrophe mechanism, and (b) whether PARGi sensitivity does indeed correlate with the relative expression levels of DNA replication genes.

## Results

### Identification of PARGi sensitive and resistant ovarian cancer cell lines

To analyse the molecular mechanisms underlying PARGi sensitivity, we assembled a panel of 10 ovarian cancer cell lines: OVMANA, Kuramochi, OVISE, RMG1, OV56, CAOV3, COV362, OVCAR3, OVSAHO and COV318 (Fig. 1A). While six of these cell lines have features typical of HGSOC, in particular *TP53* mutation (Fig. 1A), OVMANA, OVISE and RMG1 are more representative of clear cell tumours, and OV56 of low-grade serous tumours (Domcke et al., 2013; Barnes et al., 2020). Nevertheless, we reasoned that analysing a panel of sensitive and resistant cell lines would provide insight into the intrinsic cell cycle vulnerabilities responsible for PARGi sensitivity, and in particular provide insight into whether PARGi sensitivity was via a common mechanism.

**Figure 1.**
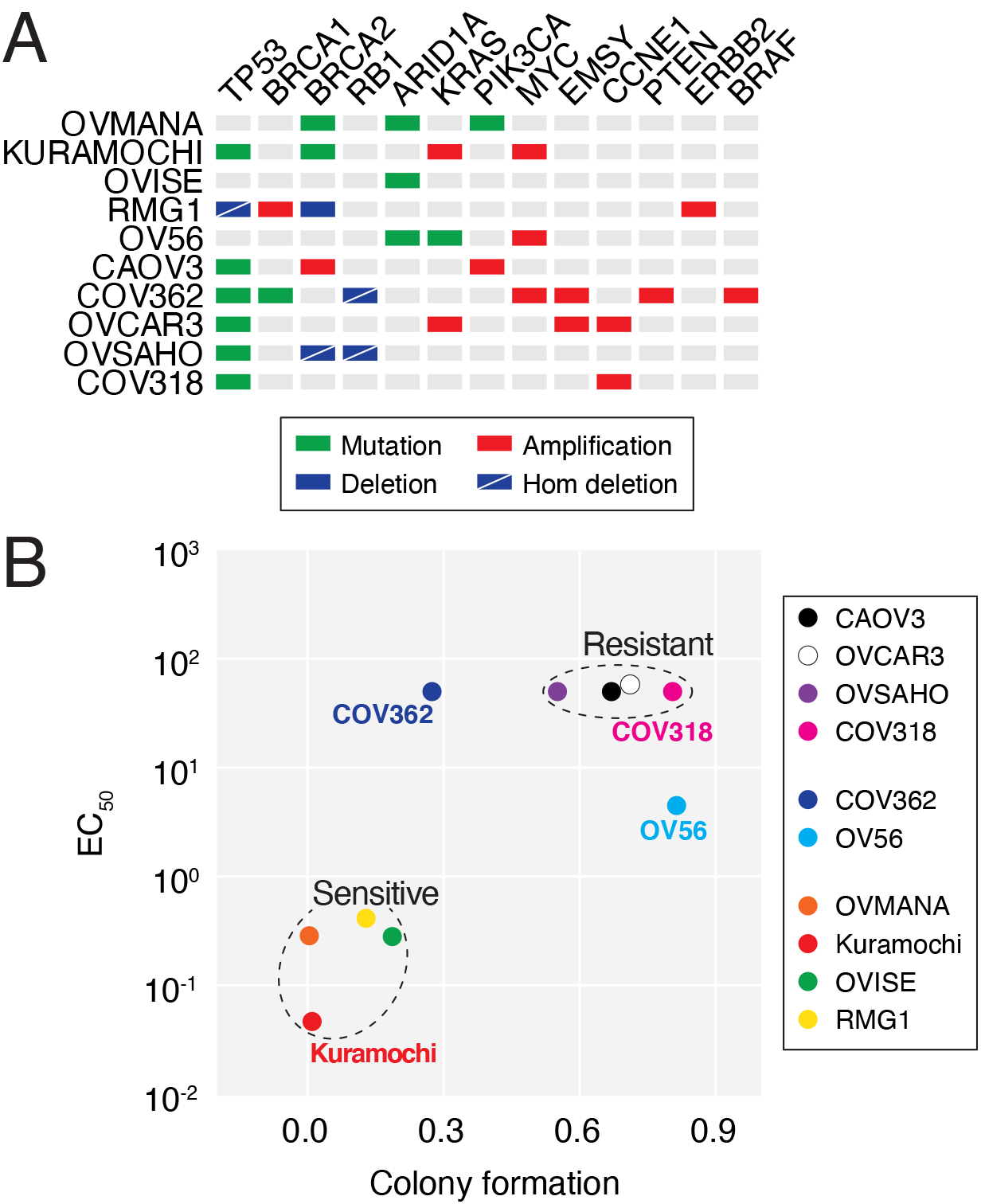
Identification of PARGi sensitive and resistant ovarian cancer cell lines. (**A**) Mutational profile of ovarian cancer cell line panel (Domcke et al., 2013). (**B**) XY plot shows correlation between EC_50_ values and colony formation with PARGi treatment. Values derived from ≥2 experiments (EC_50_) or ≥2 (colony formation). See also Figure S1.

Previously, we showed that Kuramochi, OVMANA and RMG1 are PARGi sensitive in long-term colony formation assays, while COV362, CAOV3, OVCAR3, OVSAHO and COV318 were resistant in a short-term proliferation assay. While OV56 was identified as resistant by colony formation assay, OVISE was also classified at resistant because PARGi treatment did not induce pan-nuclear γH2AX (Pillay et al., 2019). Note however that in our previous study, analysis of OVISE was challenging due to technical issues, which we have now overcome. To systematically compare PARGi sensitivity, all 10 cell lines were now analysed in long-term colony formation assays following exposure to 1 μM PDD00017273, either for 24, 48, 72 hours or continuously, and colony area quantitated. Examination of continuously-exposed cells confirmed that OVMANA, Kuramochi and RMG1 are PARGi sensitive (Fig. S1A, B). However, in contrast to our previous report, this analysis shows that OVISE is also PARGi sensitive. Outgrowth of OVMANA, Kuramochi and OVISE was also inhibited following 48- and 72-hour drug exposures (Fig. S1C), while RMG1 was not. By contrast, in line with our previous study, growth of OV56, CAOV3, OVCAR3, OVSAHO and COV318 was not markedly suppressed even in the continuous presence of PARGi, while COV362 growth was partially suppressed. In addition, two out of four PARGi-sensitive cell lines were resistant to PARPi (Kuramochi and RMG1), while four out of six PARGi resistant lines were PARPi sensitive (Fig. S1A, G, H), confirming differential sensitivity despite both inhibitors targeting PARylation.

To independently analyse PARGi sensitivity, each cell line harbouring a GFP-tagged histone was analysed by time-lapse microscopy over a 120-hour period in the presence of increasing concentrations of PARGi (Fig. S1D). Proliferation curves were then analysed to determine nuclear doubling rates and in turn EC_50_ values (Fig. S1E). COV318, OVSAHO, OVCAR3, COV362 and CAOV3 were largely unaffected even at very high concentrations of PARGi, yielding very high or indeterminant EC_50_ values (Fig. S1F). By contrast proliferation of RMG1, OVMANA and OVISE was strongly inhibited, yielding EC_50_ values in the micromolar range. Consistent with our previous analysis, Kuramochi were particularly sensitive with an EC_50_ value of 46 nM.

To integrate these two data sets, we plotted EC_50_ values from the proliferation assays against colony area in the continuous presence of PARGi (Fig. 1B). This clearly highlights OVMANA, Kuramochi, OVISE and RMG1 as PARGi-sensitive, and OVSAHO, CAOV3, OVCAR3 and COV318 as PARGi-resistant. COV362 and OV56 are more ambiguous. While OV56 appears sensitive with an EC_50_ of ∼4.5 μM, it clearly forms colonies. Closer inspection reveals that these colonies are less dense than controls (Fig. S1A), suggesting that while PARGi slows OV56 proliferation it does not block it enough to prevent outgrowth. With an indeterminant EC_50_, COV362 appears PARGi-resistant and closer inspection reveals large colonies indicating outgrowth (Fig. S1A). Thus, we conclude that both COV362 and OV56 are PARGi resistant. In summary, taking together the long-term colony formation assay and the short-term nuclear proliferation assay, we conclude that OVMANA, Kuramochi, OVISE and RMG1 are PARGi-sensitive while OV56, CAOV3, COV362, OVCAR3, OVSAHO and COV318 are PARGi-resistant.

### PARGi stabilises PAR chains in both sensitive and resistant cell lines

A possible explanation for PARGi resistance could simply be that PDD00017273 fails to engage PARG in resistant cells, for example due to drug efflux mechanisms. Therefore, we sought to determine whether PDD00017273 was indeed inhibiting PARG activity in all cases, by measuring the accumulation of PAR chains. Analysis of single cells by immunofluorescence microscopy showed that in all 10 cell lines, exposure to PARGi increased the intensity of nuclear PAR staining (Fig. 2A–C and S2A). Importantly, increased PAR staining was blocked by co-treatment with olaparib, indicating that accumulation was dependent on PARP activity. Analysis of cell populations by immunoblotting also revealed increased PAR staining in PARGi-treated cells, and importantly, this technique also showed the accumulation of higher molecular weight PAR species (Fig. 2D), indicating stabilisation of PAR chains. Interestingly, the very high molecular weight PAR species did not accumulate in OV56 or Kuramochi, and indeed, we cannot exclude that there is inter-line variation in PAR-chain dynamics. Nevertheless, we conclude that PARGi resistance is not because PDD00017273 fails to inhibit PARG activity in resistant cells, but rather because these cells tolerate the presence of stabilized PAR chains.

**Figure 2.**
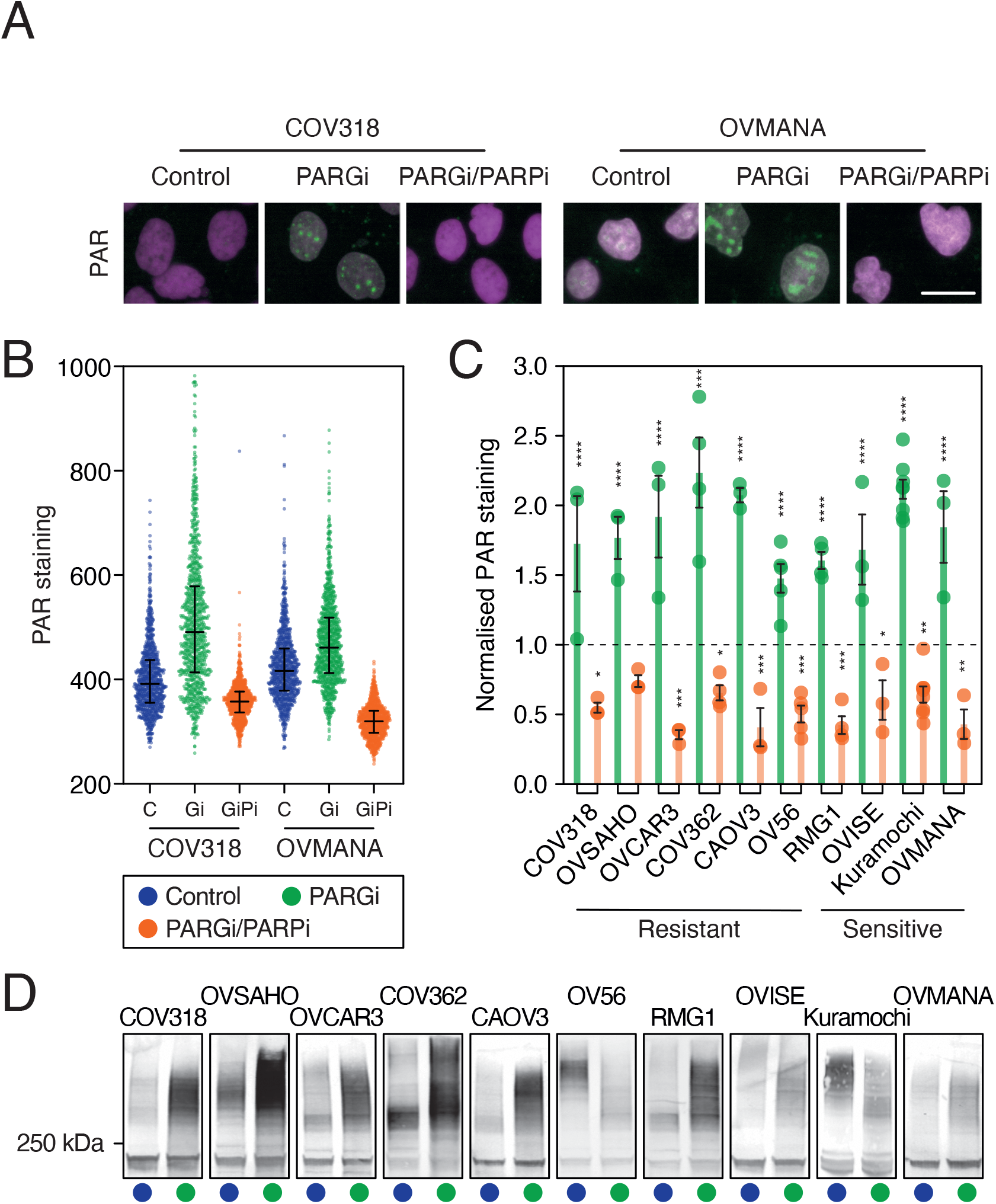
PARGi stabilises PAR chains in cell lines irrespective of PARGi sensitivity. (**A**) Representative immunofluorescence images of PAR staining in response to 48 h treatment with DMSO (Control), PARGi, and co-treatment with PARGi and PARPi in PARGi-resistant (COV318) and PARG-sensitive (OVMANA) cell lines. Scale bar: 20 μm. (**B**) Quantification of PAR staining in 1 biological replicate (dot plots, 1000 cells shown per condition). (**C**) Quantification of PAR staining in response to PARGi or co-treatment with PARGi and PARPi, normalised to DMSO. Mean of ≥3 biological replicates. (**D**) PAR was also analysed by immunoblotting. Statistics: 2-way ANOVA with Dunnett’s multiple comparisons test, selected comparisons were between drug treatments and DMSO control. Error bars represent SEM. See also Figure S2.

### PARGi sensitivity is accompanied by markers of replication stress

Having established that PAR chains accumulate with PARGi treatment in all 10 cell lines, we asked whether PARGi sensitivity occurs via a similar mechanism, in particular via DNA replication catastrophe. Previously, we showed that sensitive Kuramochi cells displayed features consistent with persistent replication stress, DNA damage and eventually replication catastrophe upon exposure to PARGi (Pillay et al., 2019). If this phenomenon is a common cause of PARGi sensitivity, we reasoned that these features should manifest in the sensitive lines but not the resistant ones. To test this, we measured accumulation of three well recognised markers of replication stress, namely γH2AX foci, RPA foci, and nuclear phospho-KAP1 (Liu et al., 2012; Toledo et al., 2017; Gralewska et al., 2020), all in single cells by immunofluorescence microscopy (Fig. 3A, B and S3A). After 48 hours exposure to PARGi, γH2AX foci were substantially and significantly increased in the sensitive lines OVISE, Kuramochi and OVMANA, but not in the six resistant cell lines (Fig. 3A and S3A). By 72 hours, cells with pan-nuclear γH2AX staining also became apparent (Fig. 3A). Induction of RPA1 foci and nuclear phospho-KAP1 showed a similar trend (Fig. 3A and S3A), confirming a correlation between PARGi sensitivity and markers of replication stress. To independently analyse replication stress, we analysed cell populations by immunoblotting to measure phosphorylation of Chk1, normalised to total Chk1, and using the ribonucleotide reductase inhibitor hydroxyurea as a positive control (Fig. 3C). Because Chk1 phosphorylation in response to HU exhibited substantial inter-line variation, we expressed the phosho-Chk1/total-Chk1 ratio as a percentage of the HU-induced maximum. This shows that PARGi induced strong Chk1 phosphorylation responses in three of the sensitive lines, RMG1, OVISE and Kuramochi, but not in four of the resistant lines, COV318, OVSAHO, OVCAR3 and COV362 (Fig. S3B), again confirming a correlation between PARGi sensitivity and replication stress.

**Figure 3.**
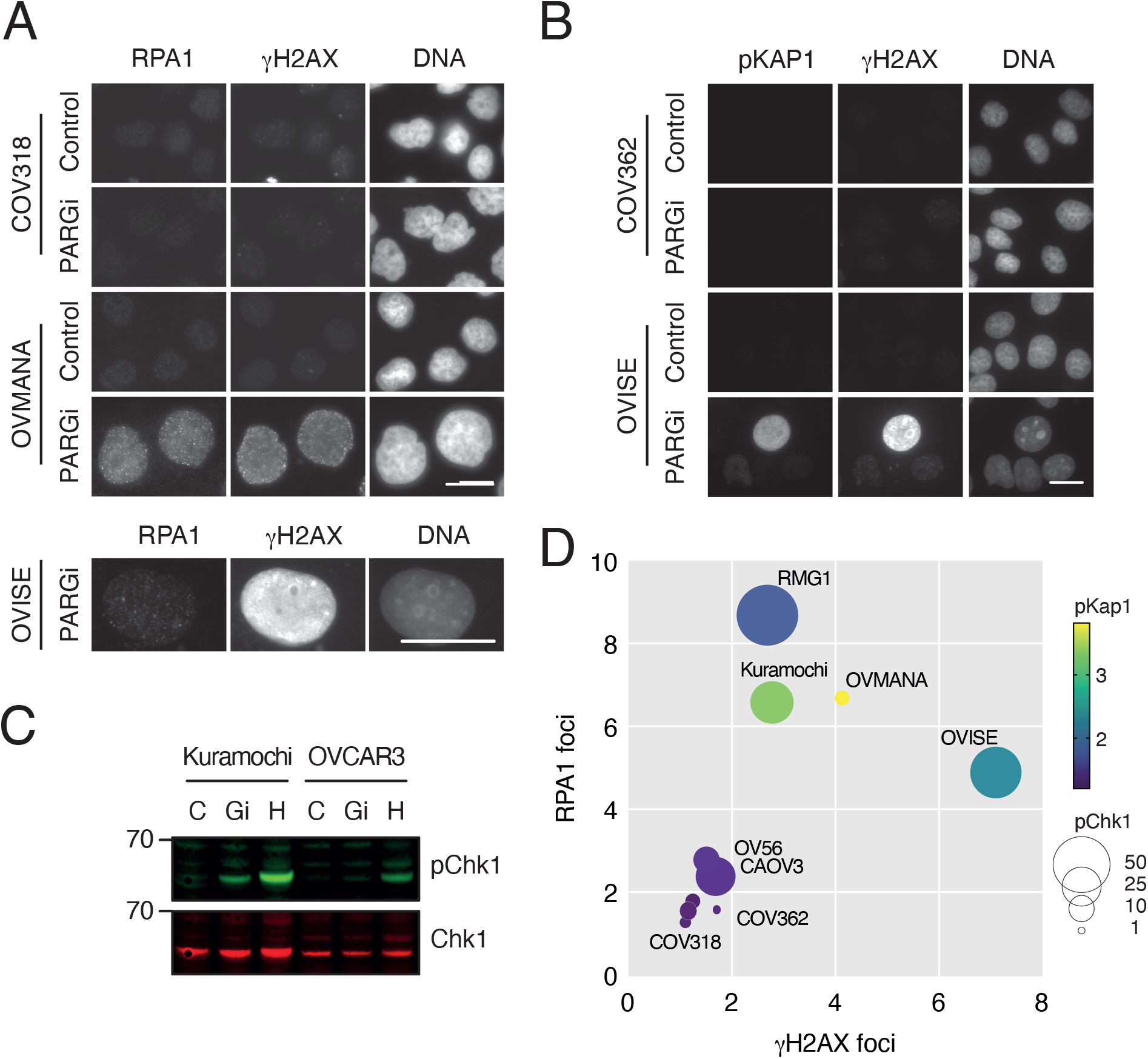
PARGi sensitivity is accompanied by markers of replication stress and the DNA damage response. **(A)**Representative of images of RPA1 and γH2AX foci in response to 48 h PARGi treatment or DMSO (control) in PARGi-resistant (COV318) and PARG-sensitive (OVMANA) cell lines (upper). RPA1 and pan-nuclear γH2AX staining also shown in PARGi sensitive OVISE cells following 72 h PARGi treatment (lower). Scale bars: 20 μm. (**B**) Representative images of pan-nuclear pKAP1 and γH2AX staining in PARGi-resistant (COV362) and PARG-sensitive (OVISE) cell lines following 72 h PARGi treatment. Scale bar: 20 μm. (**C**) Representative immunoblot Li-COR image and pChk1 quantification for PARGi-resistant (OVCAR3) and PARG-sensitive (Kuramochi) cell lines; cells were treated for 2 h with PARGi (Gi), 2mM hydroxyurea (H) as a positive control, or DMSO as a negative control (C). (**D**) Bubble plot shows fold-change in RPA1 foci, γH2AX foci, pan-nuclear pKAP1 and Chk1 phosphorylation in response to PARGi treatment. Size of bubble represents pChk1, colour of bubbles represents pKAP1. See also Figure S3.

We did notice a few exceptions to this overall trend. While RMG1 cells are sensitive and showed a substantial fold change in RPA foci and strongly induced phosho-Chk1, they showed only modest increases in γH2AX foci and phospho-KAP1. Also, while CAOV3 cells are PARGi resistant, they showed induction of RPA1 foci and Chk1 phosphorylation, but not the other markers, suggesting any replication stress resolves sufficiently to allow growth. Indeed, it has previously been reported that PARG is required for recovery from persistent, but not short-term replication stress (Illuzzi et al., 2014). These exceptions highlight the limitations of relying on only a single marker of replication stress. Therefore, we integrated these four datasets by plotting the fold changes of γH2AX foci versus RPA foci, phospho-KAP1 and phospho-Chk1 (Fig. 3D). This analysis shows a clear demarcation of sensitive and resistant cell lines, with OVISE, Kuramochi, RMG1 and OVMANA segregated from the six resistant cell lines. Thus, *in toto*, these observations demonstrate that PARGi sensitivity does reflect a common mechanism, namely persistent replication stress leading to replication catastrophe.

### PARGi induces replication fork asymmetry in sensitive cell lines

We previously showed that inhibiting PARG caused DNA replication fork asymmetry in sensitive Kuramochi cells, but not resistant OVCAR3 cells (Pillay et al., 2019), reflecting PARG’s role in restarting stalled replication forks. Having established that all four sensitive lines demonstrated features of replication stress when exposed to PARGi, we asked whether this was accompanied by persistent fork stalling, which we assessed using fork asymmetry assays. Following a 48-hour exposure to PARGi, cells were pulsed with BrdU to label active DNA replication forks, then chased with IdU to measure the speed of left and right cognate forks (Fig. 4A). While the basal level of fork asymmetry varied across the panel of 10 lines, PARGi increased fork asymmetry by 1.7–2.3 fold in three of the sensitive cell lines, OVMANA, Kuramochi and RMG1 (Fig. 4B, C). Asymmetry only increased marginally with PARGi treatment of OVISE, but notably these cells already had a high basal rate of asymmetry (Fig. 4C). By contrast, PARGi had only modest effects on the six resistant lines, with an average fold-change in fork asymmetry of 1.3. We conclude therefore that PARGi sensitivity is accompanied by an induction of DNA replication fork asymmetry, further supporting the notion that PARGi sensitivity is due to persistent replication stress.

**Figure 4.**
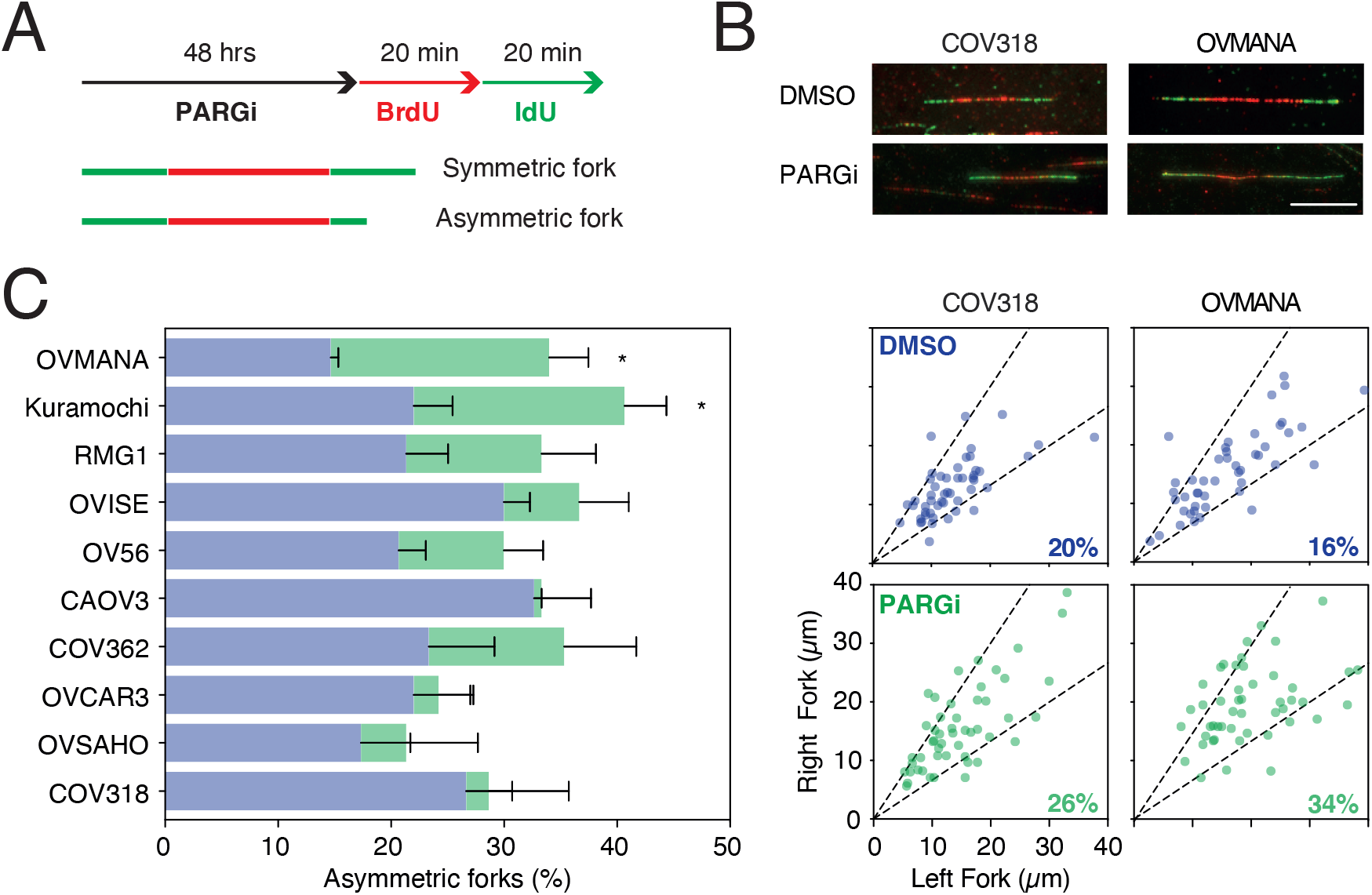
PARGi induces replication fork asymmetry in sensitive cell lines. (**A**) Schematic of DNA fibre experimental strategy. (**B**) Exemplar fibres in PARGi-resistant (COV318) and PARG-sensitive (OVMANA) cell lines. Scale bar: 10 μm. (**C**) Mean % asymmetric forks are shown on the left. The exemplar graphs on the right show correlation between R and L fork lengths. Dotted lines represent 33% cut-off, beyond which forks are considered asymmetric (% asymmetry for replicate shown in respective graph bottom right corner). Statistics: Results are from 3 biological replicates, with ≥50 forks measured per cell line, per condition. 1-way ANOVA with Sidak post-hoc test. Error bars represent SEM.

### PARGi suppresses mitotic potential in sensitive cell lines

Persistent replication stress activates intra-S and G2/M checkpoints, thereby blocking cell cycle progression, and in particular inhibiting entry into mitosis. And indeed, we previously showed that a substantial fraction of PARGi-treated Kuramochi cells underwent a Wee1-dependent G2 arrest, and that those cells that did progress through mitosis often did so abnormally, and subsequently died (Pillay et al., 2019). To determine whether a similar cell fate was shared by other sensitive lines, we analysed the panel of 10 by time-lapse microscopy for 115 hours and generated cell fate profiles (Gascoigne and Taylor, 2008). In the absence of PARGi, the vast majority of cells in each line underwent multiple successful divisions and were alive at the end of the experiment (Fig. 5A). Significantly, PARGi treatment of the sensitive cell lines resulted in an increased proportion of cells that did not enter mitosis, with 48%, 32% and 51% of OVMANA, Kuramochi and RMG1, respectively, blocking in interphase. This phenotype was not observed in the resistant cell lines where the majority of cells continued to complete multiple cell divisions when treated with PARGi.

**Figure 5.**
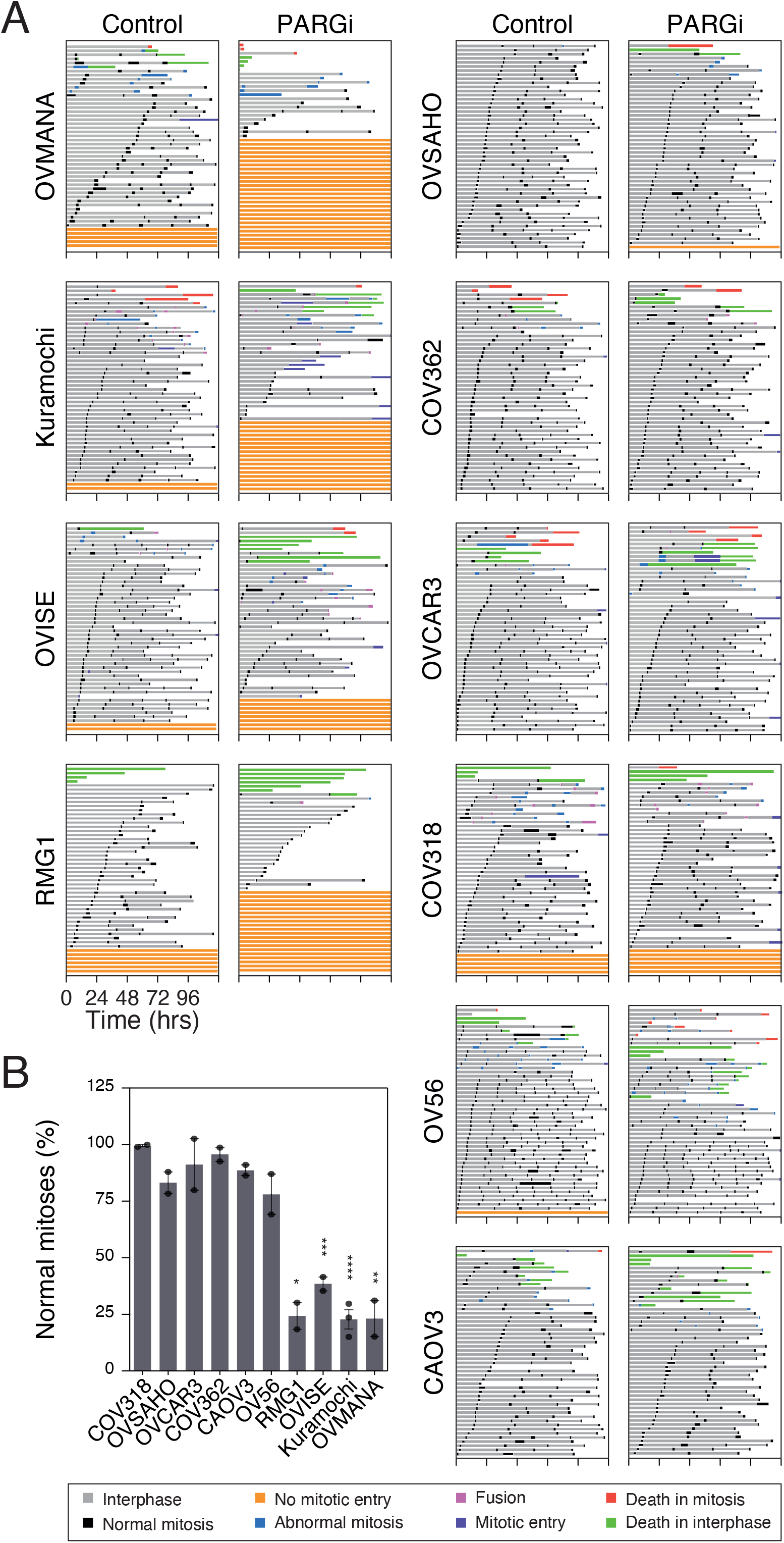
PARGi suppresses mitotic potential in sensitive cell lines. (**A**) Cell fate profiling, shows cell behaviour over 115 h. Each horizontal line represents a single cell, with the colours indicating cell behaviour. Following mitosis, one daughter cell was chosen at random to continue the analysis. No mitotic entry does not include death in interphase where mitosis does not take place. Abnormal mitosis includes cell division, abnormalities such as tripolar cell divisions, binuclear daughter cells and division of binuclear cells. Slippage is recorded where cells enter mitosis, then exit without division. Fusion was recorded where daughter cells appeared to separate but subsequently joined back together. (**B**) Graph shows the % of total normal mitoses in PARGi-treated compared to DMSO controls. Represents means of ≥2 biological replicates. Statistics: total normal mitoses in DMSO-treated compared to PARGi-treated cells was compared by 1-way ANOVA with Sidak post-hoc test. Error bars represent SEM.

Interestingly, in the four sensitive cell lines, a small proportion of cells did not enter mitosis in the absence of PARGi, consistent with these cells exhibiting an underlying vulnerability that is markedly exacerbated by inhibition of PARGi. Of the resistant lines, a small proportion of COV318 cells also did not enter mitosis in the absence of PARGi, but in contrast to the sensitive lines, this did not increase substantially upon exposure to PARGi.

In most of the cultures analysed, a fraction of cells underwent abnormal mitoses and/or cell death and this phenotype was exacerbated by PARGi in sensitive, but not resistant lines. In particular, while only 14% of PARGi-treated OVISE cells blocked in interphase, the number of cells undergoing abnormal mitoses and/or cell death increased from 17% to 46%. Although the analyses above indicate that OV56 is PARGi resistant, the cell fate profiling indicates that drug exposure increases the number of abnormal mitoses from 23% to 35% (Fig. 5A), possibly reflecting a synthetic effect with the prolonged cell culture and/or imaging conditions.

To quantitate these PARGi effects on mitotic potential, we counted the number of productive cell divisions in control and drug-treated populations. In sensitive cell lines, PARGi dramatically reduced the number of normal mitoses, by an average of 73%, compared with a minor reduction, (11% on average), in resistant lines (Fig. 5B). Thus, considering both the cell fate profiles and the quantitation of successful divisions, we conclude that sensitivity to PARGi is accompanied by a dramatic loss of mitotic potential, in particular a pre-mitotic block, consistent with activation of intra-S and G2/M checkpoints due to persistent replication stress.

### PARGi sensitivity correlates with replication stress

The analyses described above show that markers of replication stress (Fig. 3), fork asymmetry (Fig. 4) and pre-mitotic block (Fig. 5) do appear to correlate with PARGi sensitivity. To confirm this, we integrated the various datasets by scaling each parameter so that the most resistant and sensitive lines scored 1 and 0 respectively (Fig. 6A). These were then averaged to yield an overall sensitivity score and rank ordered. Importantly, the various cell biological parameters clearly align with sensitivity as defined by the colony formation and short-term proliferation assays, with COV318, OVSAHO, OVCAR3 and COV362 ranking as the most resistant lines, and OVMANA, Kuramochi and OVISE the most sensitive. Of the sensitive lines, RMG1 ranks as the least sensitive and indeed, as noted above, this line requires continuous exposure for fully penetrant inhibition (Fig. S1A–C). Of the resistant lines, OV56 is the least resistant and, again as noted above, although colonies form in the presence of PARGi, they are less dense than control colonies (Fig. S1A–B). CAOV3 are also at the more ‘sensitive’ end of the resistant spectrum and interestingly, these cells do display increased RPA foci when exposed to PARGi (Fig. S3A), suggesting that while PARGi does induce replication stress, it is resolved sufficiently to allow efficient recovery. Nevertheless, this integrated analysis confirms that PARGi sensitivity does indeed correlate with markers of replication stress, suggesting a common underlying mechanism. Moreover, because PAR chains are stabilized in all cell lines analysed (Fig. 2C, D), an important corollary is that resistant cell lines can efficiently complete DNA replication despite the presence of stabilized PAR chains.

**Figure 6.**
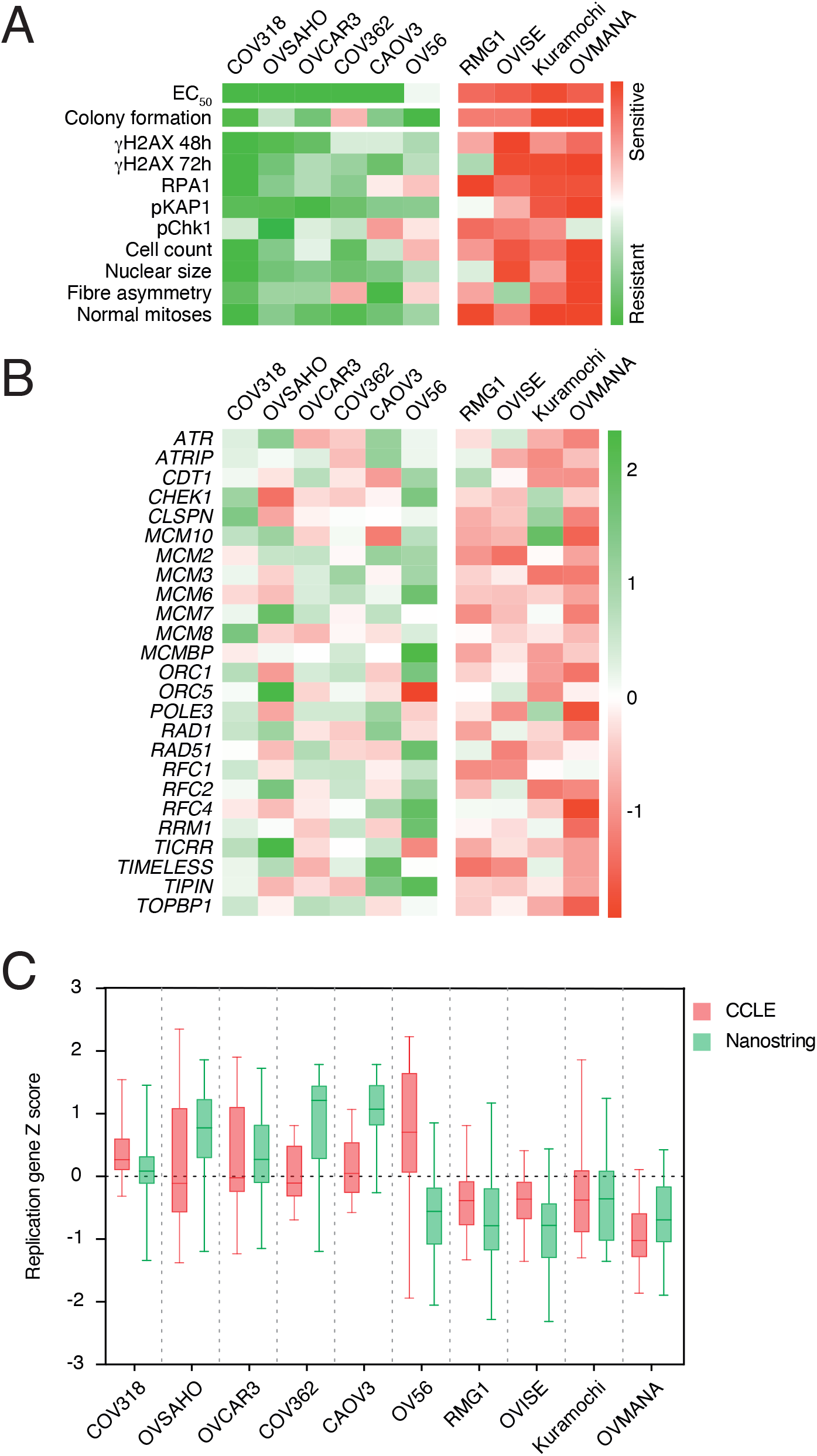
PARGi-sensitive cell lines have lower expression of replication genes. **(A)** PARGi-sensitivity assay results summarised as a heatmap. Assay results (except EC_50_s) were scaled from 0–1, averaged, and cell lines rank ordered accordingly. (**B**) NanoString replication gene Z scores represented as a heatmap. (**C**) Box-whisker plots (min–max) for replication genes Z scores shown for NanoString compared with CCLE.

### PARGi sensitivity correlates with lower expression of DNA replication genes

The strong correlation between markers of replication stress and PARGi sensitivity supports our previous conclusion that sensitive cell lines harbour an underlying DNA replication vulnerability that makes them particularly dependent on PARG activity to re-start stalled replication forks (Pillay et al., 2019). While the nature of this vulnerability remains to be determined, when we previously interrogated CCLE microarray data, we noted that in addition to Kuramochi, the expression levels of a number of DNA replication genes was lower in RMG1, OV56, OVMANA and OVISE cell lines compared with resistant lines (Pillay et al., 2019). Of these four, we previously showed that RMG1 and OVMANA are PARGi sensitive, and here we demonstrate that OVISE is also sensitive. However, while OV56 is the most “*sensitive*” of the resistant lines, it is resistant despite apparent low expression levels of DNA replication genes. To address this anomaly, we set out to independently validate the CCLE data by analysing the expression levels of 25 DNA replication genes using a custom NanoString CodeSet (Fig. 6B). This analysis confirms that these DNA replication genes are down-regulated in sensitive cell lines compared with resistant lines. However, in contrast to the CCLE data, our data shows that, like the other resistant lines, the DNA replication genes are over expressed in OV56 (Fig. 6C). Interestingly, while RNAseq data released more recently by the CCLE project is consistent with their previous microarray data (Barretina et al., 2012; Ghandi et al., 2019), an independent RNAseq study indicates that DNA replication genes are up-regulated in OV56. In brief, we analysed read counts of these 25 replication genes in an independent RNAseq analysis of 41 epithelial ovarian cancer cell lines (Klijn et al., 2015) (GEO accession number GSE40788). This revealed higher expression of these genes in OV56 cells than indicated in the CCLE data set, in agreement with our NanoString data (data not shown). Aside from this one anomaly, two independent RNAseq datasets (Klijn et al., 2015; Ghandi et al., 2019), and our NanoString-based analysis, indicate that cell lines that are sensitive to PARGi tend to have lower expression levels for a number of DNA replication genes. Thus, taking together the drug-sensitivity data, the markers of replication stress and the gene expression studies, we conclude that pharmacological PARG inhibition induces replication catastrophe in ovarian cancer cells with down-regulated DNA replication genes.

### PARPi versus PARGi sensitivity

In terms of PARPi sensitivity, using a short-term confluence assay, we previously showed that OVSAHO, COV318, COV362 and CAOV3 were PARPi resistant, whereas OVCAR3 was sensitive (Pillay et al., 2019). Because long-term target engagement can be required to observe synthetic lethality between cells with *BRCA* mutation and olaparib (James et al., 2016), we re-visited this and compared PARPi and PARGi sensitivity in long-term colony formation assays with continuous drug exposure. Of these four lines, with the exception of COV318, all showed a reduction in outgrowth in response to PARPi (Fig. S1G). By contrast, and in agreement with our previous report, OV56, Kuramochi and RMG1 were relatively resistant to PARPi. Notably, OVISE and OVMANA were sensitive to both PARPi and PARGi.

Thus, we conclude that of the ten ovarian cancer cell lines analysed here, four are PARPi-sensitive and PARGi-resistant; two are PARGi-sensitive and PARPi-resistant; two are sensitive to both modalities and two are resistant to both (Fig. S1H). This implies that sensitivity to PARGi and PARPi is neither mutually exclusive nor overlapping, indicating that the determents of sensitivity are complex and context dependent. In this regard OV56 is an interesting case; while categorized as PARGi-resistant, it is the least resistant of the resistant lines and PARGi exposure is clearly not completely inconsequential; the short-term proliferation assay yields an EC_50_ value, albeit a high one, and the colonies are less dense than controls for in the PARGi-treated culture. And yet in response to PARPi, it forms robust colonies. Thus, despite the intimate relationship between PARP and PARG activities, and despite the complexities described above, a differential sensitivity clearly manifests in a number of cell lines, indicating that PARG inhibitors may offer an alternative therapeutic option to target a subset of ovarian cancers not expected to be PARPi-sensitive.

## Discussion

We set out to address two questions; (1) whether PARGi-sensitive ovarian cancer cells exhibit similar or different phenotypes in response to PARG inhibition, and (2) whether PARGi sensitive cells exhibit reduced levels of DNA replication genes. Our analysis of four PARGi-sensitive and six resistant cell lines revealed that all the sensitive cell lines displayed features of persistent replication stress, DNA damage and eventually replication catastrophe, evidenced by accumulation of asymmetric DNA replication fibres, γH2AX and RPA foci, KAP1 and Chk1 phosphorylation, and, following prolonged exposure, a pan-nuclear γH2AX phenotype that indicates RPA exhaustion. While synthesising our observations shows that these replication stress markers track with PARGi sensitivity, we did observe some idiosyncratic responses. For example, while RMG1 cells are PARGi sensitive and underwent a pre-mitotic block with PARGi treatment, they displayed only modest induction of γH2AX foci and KAP1 phosphorylation. Conversely, PARGi-resistant CAOV3 showed a significant induction of RPA foci. These idiosyncrasies, which demonstrate the importance of analysing multiple markers, could reflect both technical and biological issues. For example, for technical purposes all 10 cell lines were subjected to the same experimental workflows to allow direct comparison, despite differences in doubling times, growth dynamics and EC_50_ values. In terms of biological differences, PARGi resistance could conceptually arise either because drug-treated cells do not accumulate replication stress, or because they are better able to resolve and recover from problems with DNA replication. This distinction may explain why COV318 and OVSAHO for example are particularly PARGi-resistant, while CAOV3 display some features of replication stress, and although OV56 is categorised as resistant, exposure to PARGi is not benign.

A key issue is whether PARG inhibitors will offer distinct therapeutic opportunities compared with PARP inhibitors in the treatment of HGSOC. Of the six PARGi-resistant cells, four are sensitive to olaparib. Previously we noted that OVSAHO, COV362 and CAOV3 were PARPi-resistant (Pillay et al., 2019). This was based on a short-term confluency-based proliferation assay, whereas here we performed long-term colony formation assays and subsequently found these three cell lines to actually be PARPi-sensitive. This issue highlights the need for orthogonal approaches to determine sensitivity and here for PARGi we employed both short-term and long-term assays. It is also worth noting that to monitor the effects of PARGi in short-term proliferation assays, we measured nuclear doubling by tracking GFP-H2B nuclei rather than confluency, which can be confounded by cells increasing in size despite not proliferating. This is particularly important in the context of persistent replication stress where cells can adopt a reversible senescence-like phenotype, characterised by large, flat cellular morphology (Maya-Mendoza et al., 2014; Maya-Mendoza et al., 2016). Therefore, of the four PARGi-sensitive cell lies, only two are sensitive to olaparib, consistent with previous reports that PARGi and PARPi sensitivity are often non-overlapping (James et al., 2016; Pillay et al., 2019).

This non-overlapping sensitivity to PARPi and PARGi may seem counter-intuitive, as PARP1/2 and PARG activities work in concert to repair DNA damage. One might expect therefore that like PARPi, PARGi would also be toxic towards tumour cells with DDR defects (Chen and Yu, 2019). Indeed, some studies suggest synthetic lethality between *BRCA1/2* mutations and loss of PARG (Gravells et al., 2017; Chen and Yu, 2019). However other studies do not (Noll et al., 2016; Pillay et al., 2019). Moreover, two of the PARGi-resistant lines, COV362 and OVSAHO, have reported *BRCA* defects (Domcke et al., 2013), and are PARPi-sensitive, indicating that *BRCA* mutation does not predict PARGi sensitivity. By contrast, our observations suggest that PARGi sensitivity is better predicted by interrogating expression levels of DNA replication genes. Interestingly, the two lines sensitive to both PARGi and PARPi, namely OVISE and OVMANA are reported to be *TP53* wildtype. Whether these cells lines are p53-proficient is unclear, however, because *TP53* is near-ubiquitously mutated in HGSOC, the PARGi/PARPi differential may be more marked in this setting. To test this, we are evaluating PARPi and PARGi sensitivity in an array of patient-derived HGSOC samples (Nelson et al., 2020), and thus far our analysis shows that in general PARGi-sensitive *ex vivo* cultures are not PARPi sensitive and vice versa (Morgan, Nelson and Taylor, unpublished).

The second key aim of this study was to ascertain whether PARGi sensitive cells exhibit reduced levels of DNA replication genes. Interrogating CCLE microarray data, previously identified four cell lines which were of interest in that, like Kuramochi, they displayed reduced expression of DNA replication genes. Of these four, we showed two to be PARGi sensitive (Pillay et al., 2019). Here we demonstrate that a third, OVISE, is also sensitive and that DNA replication genes are in fact not down-regulated in the fourth cell line, OV56. Thus, based on our analysis of 10 cell lines, there is indeed a strong correlation between PARGi sensitivity and the expression levels of a subset of DNA replication genes. Moreover, interrogating DNA replication genes as a means to identify PARGi sensitive lines performed better than originally reported and is thus encouraging in terms of developing a predictive biomarker of PARGi sensitivity. Why DNA replication genes are down-regulated in a subset of ovarian cancer cell lines remains to be determined. Furthermore, whether these genes are “*down-regulated*”, e.g. due to active suppression of gene expression, is not clear. Indeed, an alternative possibility is that resistant cell lines have actively up-regulated DNA replication genes. Distinguishing between these two possibilities will require further analysis but it has previously been shown that DNA replication genes can be up-regulated as an adaptive response to replication stress. Indeed, replication stress reduces cell fitness of cancer cells, and several adaptive mechanisms have been described including upregulation of translesion synthesis (TLS) or replisome components (Bianco et al., 2019; Nayak et al., 2020). Bianco *et al*, reported over-expression of Timeless, Claspin and Chk1 in various cancers, and showed that tolerating oncogene-induced replication stress was accompanied by overexpression of DDR genes and genes involved in fork protection. Interestingly, these overexpressed genes included 9 of the 25 included in our “*replication stress*” signature, including Claspin and Timeless (Fig. 6). Thus, differential PARGi sensitivity might reflect different adaptive responses to buffer oncogene-induced DNA replication stress. If some tumours adapt by up-regulating DNA replication genes, this would engender intrinsic PARGi resistance. However, if other tumours buffer replication stress via different routes, i.e. those that do not require upregulation of DNA replication genes, this may render them intrinsically PARGi sensitive. Note that parallel evolutionary trajectories to increase fitness in response to replication stress have been described in yeast (Fumasoni and Murray, 2020). If this is the case, one might expect that PARGi-sensitive cells could acquire resistance via up-regulating DNA replication genes, a prediction that could be tested *in vitro*. Also, if intrinsic PARGi-resistance does indeed reflect an adaptive response that leads to activating a gene expression programme that converges on DNA replication genes, understanding the signalling pathways and transcription factor programmes involved could open up further opportunities to develop biomarkers of PARGi-resistance, in turn aiding the identification of tumours that are more likely to be PARGi-sensitive.

PARG inhibitors suitable for *in vivo* testing and clinical evaluation have not yet been described. However, our observations may aid the design of first-in-human trials testing clinical candidates when they become available. In light of the excellent efficacy of PARP inhibitors in HRD tumours, efforts to evaluate PARG inhibitors in patients will focus on tumours likely to be PARP inhibitor resistant. Indeed, because 50% of HGSOC cases are predicted to be HR-proficient, there is a pressing need to develop therapeutic strategies to benefit women unlikely to benefit from PARP inhibitors. However, it is unlikely that all PARPi-resistant tumours will respond to a PARG inhibitor. Therefore, critical to the design of trials testing PARG inhibitors will be development of suitable enrichment biomarker that predicts sensitivity. Here we show that PARGi sensitivity is underpinned by a common mechanism, namely persistent replication stress. Moreover, we show that PARGi-sensitive cell lines exhibit reduced levels of DNA replication genes, suggesting that a ‘‘replication stress” signature may have merit as a predictive biomarker. Important next steps will be to evaluate these genes in a wider collection of patient-derived HGSOC samples with defined PARGi sensitivity.

## Experimental Procedures

### Materials

Small molecule inhibitors were dissolved in DMSO and used at the following concentrations, unless stated otherwise: PARG inhibitor, PDD00017272 (PARGi), 1 μM (James et al., 2016); PARP1/2 inhibitor, Olaparib (PARPi), 1 μM (Selleckchem). Hydroxyurea (Sigma) was dissolved in water and used at 2 mM. BrdU and IdU (Sigma) were dissolved in culture media and used at concentrations described for DNA fibre assay below.

### Human Cell Lines

The human ovarian cancer cell lines OVCAR3 (ATCC), Kuramochi, OVMANA, OVSAHO, OVISE (JCRB Cell Bank) were grown in RPMI; COV362, COV318 (Sigma) and CAOV3 (ATCC) were grown in DMEM; RMG1 (JCRB Cell Bank) was grown in Ham’s F12; all supplemented with 10% foetal bovine serum, 100 U/ml streptomycin, 100 U/ml penicillin, 2 mM glutamine and maintained at 37°C in a humidified 5% CO2 atmosphere. OV56 (Sigma) was grown in DMEM/F12 with foetal bovine serum reduced to 5% and supplemented with 10 ug/ml insulin, 0.5 ug/ml hydrocortisone, 100 U/ml streptomycin, 100 U/ml penicillin, 2 mM glutamine, and maintained as above. All lines were authenticated by the Molecular Biology Core Facility at the CRUK Manchester Institute using Promega Powerplex 21 System, and underwent periodic testing for mycoplasma.

### Colony formation assay

1000 cells per well were seeded into 6-well plates 24 h prior to drug treatment. DMSO/PARGi/PARPi was added; cells were either treated continuously or the drugs were washed out at the specific time points as indicated in the figures. After colonies had developed (range: 2–4 weeks, depending on cell line), cells were fixed in 1% formaldehyde and stained for 10 min with 0.05% (w/v) crystal violet solution (Sigma Aldrich). Colonies were then rinsed with ddH2O. Plates were then imaged using a ChemiDoc™ Touch Imaging System (BioRad), and analysed with an ImageJ ‘colony area’ plug-in developed by (Guzmán et al., 2014).

### Immunofluorescence

Cells were plated onto 13 mm coverslips 24 h prior to drug treatment. For the RMG1 cell line, coverslips were coated with 0.01% Poly-L-Lysine (Sigma Aldrich). Cells were washed and fixed in 1% formaldehyde, quenched in glycine, then incubated with primary antibodies (mouse anti-PAR, 1:400 Merck Millipore Cat#AM80 RRID: AB_2155072; mouse anti-γH2AX pS139, 1:2000 Merck Millipore Cat#05-636 RRID: AB_309864; rabbit anti-pKAP1, 1:500 Bethyl Laboratories Cat#A300-767A RRID: AB_669740; rabbit anti-RPA70, 1:500 Abcam Cat#ab79398 RRID: AB_1603759) in PBS-T (PBS, 0.1% Triton X-100); or 1% dried skimmed milk (Marvel) in PBS-T after 15 min blocking in 1% milk for PAR staining) for 1 h at room temperature. Coverslips were washed x3 in PBS-T and incubated with the appropriate fluorescently conjugated secondary antibodies (1:500; Donkey anti-rabbit Cy2, Cat#711-225-152 RRID: AB_2340612; Donkey anti-rabbit Cy3, Cat#711-165-152 RRID: AB_2307443; Donkey anti-mouse Cy2, Cat715-225-150 RRID: AB_2340826; Donkey anti-mouse Cy3, Cat#715-165-150 RRID: AB_2340813 all Jackson ImmunoResearch Laboratories Inc) for 30 min at room temperature. Coverslips were washed 3x in PBS-T and DNA stained for 1 min with 1 μg/ml Hoechst 33258 (Sigma Aldrich) at room temperature. Coverslips were washed 3x in PBS-T and mounted (90% glycerol, 20 mM Tris, pH 9.2) onto slides. Image acquisition was using an Axioskop2 (Zeiss, Inc.) microscope fitted with a CoolSNAP HQ camera (Photometrics) using MetaMorph Software (Molecular Devices). For high-throughput immunofluorescence, cells were processed as above in a 96-well plate format (PerkinElmer Cell Carrier plates), with two additional final washes in PBS. Images were acquired using Operetta^®^ High Content Imaging System (Perkin Elmer), and quantified using Columbus High Content Imaging and Analysis Software (Perkin Elmer). Mean fluorescence intensity within the nuclear area, or foci quantification within the nuclear area using Columbus ‘spot finder’ tool, was quantified as a mean value per cell. These were also calculated for a secondary antibody only control. For final values, secondary antibody only control values were subtracted from the values from the stained cells with equivalent treatment.

### Drug sensitivity assay and cell fate profiling

Cells were seeded at 500–8000 cells per well in a 96-well plate (Greiner Bio-One/ PerkinElmer Cell Carrier), 24 h prior to drug treatment. For EC_50_s cells expressing GFP-H2B were used and PARGi was serially diluted from 100 μM–0.381 nM (19 concentrations in total). Following drug treatment, cells were imaged using an IncuCyte^®^ ZOOM (Essen BioScience) equipped with a 20X objective and maintained at 37°C in a humidified 5% CO_2_ atmosphere. Nine phase contrast and fluorescence images per well (for GFP-H2B expressing cells) were collected every 6 h for a total of 120 h when analysing proliferation and drug sensitivity or every 10 min for a total of 115 h for cell fate profiling. IncuCyte® ZOOM software was used in real-time to measure confluence and green object count. Green object count was then used to generate dose-response curves in PRISM, from which EC_50_ values were calculated. In highly resistant cell lines, EC_50_ could not be determined accurately, and it was assigned an approximate value of 50 μM (half the maximum dose tested). For cell fate profiling, image sequences were exported in MPEG-4 format and analysed manually to time and annotate cell behaviours. Prism 8 (GraphPad) was used for statistical analysis and presentation.

### Immunoblotting

Cells were treated with DMSO/PARGi for 48 h prior to harvesting, or for 2 h with 2 mM HU as a positive control. Proteins were extracted, quantified by Bradford assay, then denatured by boiling in sample buffer (0.35 M Tris pH 6.8, 0.1 g/ml sodium dodecyl sulphate, 93 mg/ml dithiothreitol, 30% glycerol, 50 μg/ml bromophenol blue). Proteins were then resolved by SDS-PAGE, then electroblotted onto Immobilon-Fl PVDF membrane (Millipore). In place of a loading control, REVERT total protein stain solution (LI-COR) was used for normalisation. The membrane was incubated with REVERT solution for 5 min, followed by washing in 6.7% (v/v) Glacial Acetic Acid, 30% (v/v) Methanol in water. Before imaging on the Odyssey ® CLx Imaging System (Li-COR), the membrane was washed with water. Following imaging, REVERT stain was removed, using REVERT reversal solution (0.1 M NaOH, 30% v/v methanol in water). Membranes were then blocked using 5% dried skimmed milk (Marvel; or 5% BSA for pChk1) TBS-T (50 mM Tris pH 7.6, 150 mM NaCl, 0.1% Tween-20). Primary antibodies (mouse anti-Chk1, 1:500 Santa Cruz Biotechnology Cat#sc-8408 RRID: AB_627657; rabbit anti-pChk1, 1:750, Cell Signalling Cat#2348 RRID: AB_331212; mouse anti-PAR, 1:400 Merck Millipore Cat#AM80 RRID: AB_2155072;) were diluted in 5% dried skimmed milk (Marvel), 0.2% Tween-20, 0.01% SDS TBS, or 5% BSA 0.2% Tween-20, 0.01% SDS TBS (pChk1). Membranes were then washed three times in TBS-T and incubated for 1 h with appropriate fluorescently-conjugated secondary antibodies (1:5000, IRDye ® 800CW donkey anti-rabbit Cat#925-32213 RRID: AB_2715510; IRDye ® 680RD donkey anti-mouse, Cat#926-68072 RRID: AB_10953628; both LI-COR) in 5% dried skimmed milk (Marvel) 0.2% Tween-20 + 0.01% SDS TBS. Membrane was rinsed with TBS, before imaging on Odyssey ® CLx Imaging System (Li-COR).

### Lentiviral production and Transduction

To produce the GFP-H2B cell lines, AAV293T cells were plated at 5 × 10^4^ cells per well in a 24-well plate. Media was replenished 1 h before transfection. Cells were transfected with pLVX-based lentiviral plasmids (Takara Bio), modified to express human histone H2B tagged at the N-terminus with GFP (pLVX-myc-EmGFP-H2B) plus psPAX2 and pMD2.G (gifts from Didier Trono, Addgene) using 16.6 mM CaCl_2_ in DMEM supplemented with 10% Hyclone™ serum (GE Healthcare) and incubated overnight. Virus was harvested 48 h after transfection, centrifuged and filtered (0.45 μm). Cells were seeded at 2–10 × 10^5^ cells per well in a 12-well plate and diluted lentivirus and 10 μg/ml polybrene added 48 h later. The plates were centrifuged at 300xg at 30°C for 2.5 h. 1 ml of culture media was added and the plates incubated overnight. Puromycin (1 μg/ml, 2 μg/ml for OVMANA) was added 48 h post-transduction.

### DNA fibre assay

#### Sample preparation

Sub-confluent cells were incubated in the presence of PARGi/DMSO for 48 h, then pulsed with BrdU at 5 μM plus PARGi/DMSO for 20 min. This was followed by 3 washes with warm PBS, pulsing with 200 μM IdU (Sigma) plus PARGi/DMSO for a further 20 min, then washing twice with ice-cold PBS. Following trypsinization, cells were diluted in ice-cold PBS to give a final concentration of 1–5 × 10^5^ cells/ ml and kept on ice.

#### Slide preparation

The cell suspension (2 μl) was then dropped onto microscope slides and dried at room temperature for 5–10 min before mixing with 7 μl of spreading buffer (200 mM Tris-HCl pH 7.5, 50 mM EDTA, and 0.5% SDS), and incubating for a further 5 min. Slides were tilted approximately 5–10° so that the cell suspension runs across the length of the slide. Slides were air dried and fixed in methanol/acetic acid (3:1) for 10 min, air dried and stored at 4°C.

#### Immunostaining

Prior to immunostaining, slides were washed twice with H2O for 5 min, 1 × 2.5 M HCl, denatured with 2.5 M HCl for 1 h, rinsed twice with PBS and then washed with blocking solution (PBS with 1% BSA and 0.1% Tween-20) twice for 5 min and then for 1 h. For immuno-labelling all antibodies were dissolved in blocking solution. Slides were then incubated with a rat anti-BrdU antibody (BU1/75 [ICR1] 1:500 Abcam Cat# 6326; RRID: AB_305426) to detect BrdU for 1 h under humidified conditions, rinsed 3 × PBS, fixed for 10 min with 1% formaldehyde, rinsed 3 × PBS, and quenched in glycine. Slides were then rinsed 3 × PBS followed by overnight, 4°C incubation with mouse anti-BrdU (B44, 1:100 BD Biosciences Cat# 347580; RRID: AB_400326) to detect IdU. Slides were then washed twice with PBS, 3 times for 5 min in blocking solution, followed by incubation in the appropriate fluorescently conjugated secondary antibodies diluted in blocking solution (1:500; Donkey anti-rat Cy3 Cat# 712-165-153; RRID: AB_2340667; Donkey anti-mouse Cy2, Cat715-225-150 RRID: AB_2340826; all Jackson ImmunoResearch Laboratories Inc) for 1.5 h. Post-incubation, slides were washed 2 × PBS, 3 × 5 min with blocking solution and 2 × PBS. All slides were mounted to coverslips using PBS/Glycerol (1:1). *Imaging and quantitation*: Images were acquired using an Axioskop2 (Zeiss, Inc.) microscope fitted with a CoolSNAP HQ camera (Photometrics) and 2–5 slides analysed per condition. Fibre lengths were quantified using ImageJ software (NIH).

### RNA extraction

RNA extraction was performed on pellets from sub-confluent cells scraped from cell culture flasks, using RNeasy kit (QIAGEN), according to the manufacturer’s instructions. RNA concentration was measured using Nanodrop.

### NanoString

100 ng RNA was provided at concentration of 4 ng/μl to the genomic technology facility, who performed the NanoString experiment. Briefly: RNA was hybridized with custom nCounter Reporter and Capture probe sets (custom CodeSet of 29 genes (NanoString Technologies), consisting of 25 replication genes (see Fig. 6 for list) and 4 reference genes previously reported to be appropriate for use as normalisation controls in ovarian cancer (RPLP0, PPIA, IPO8, TBP) at 65°C overnight, unhybridized probes removed, complexes bound to the imaging surface, and images acquired using the nCounter Digital Analyzer. Data was analysed using nSolver Analysis software 4.0. QC, which generated normalised transcript counts.

### Bioinformatics

Read counts of the same 25 replication genes were analysed in an independent RNAseq analysis of 41 epithelial ovarian cancer cell lines provided by (Klijn et al., 2015) (GEO accession number GSE40788).

### Quantification and Statistical Analysis

Statistical analysis was carried out using GraphPad Prism 8 Software. P values were designated as follows: * <0.05, ** <0.01, *** <0.001, **** <0.0001, ns p>0.05. Details of statistical analyses are described in the figure legends.

## Acknowledgments

We thank the members of the Taylor lab for advice and comments on the manuscript and the CRUK MI for core facilities. The research was funded by Cancer Research UK Programme Grant to S.S.T (C1422/A19842).

## Author contributions

Methodology, Investigation, Validation and Formal Analysis, C.C-G., R.D.M., L.N. and B.B. Writing C.C-G., J.M. and S.S.T. Conceptualisation, Funding and Supervision, S.S.T.

## Supplementary Figure legends

**Figure S1.**
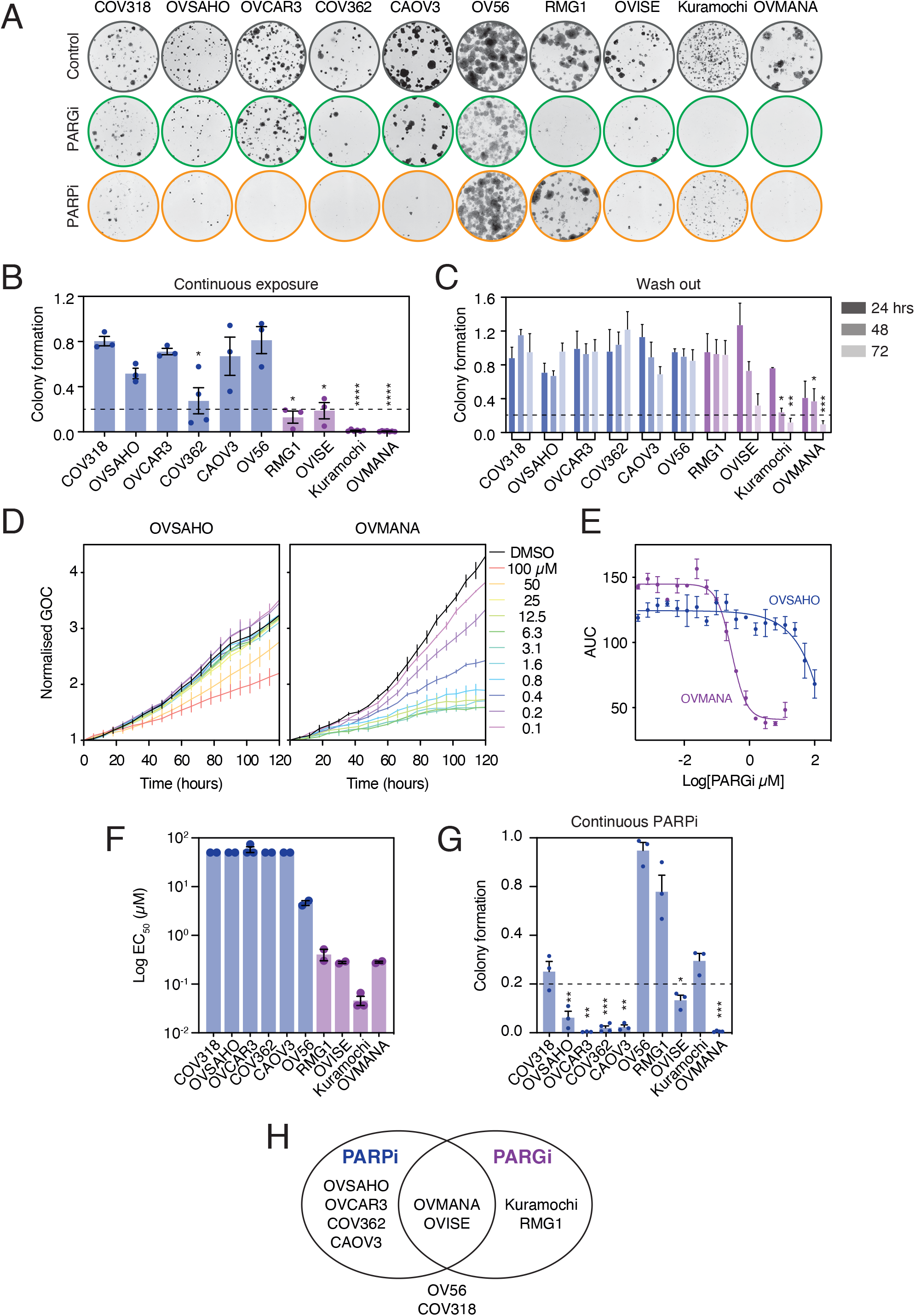
Ovarian cancer cell lines exhibit differential sensitivity to PARGi and PARPi. (**A**) Colony formation in the continuous presence of PARGi or PARPi. Representative of ≥3 biological replicates. (**B**) Quantification of colony assays with constant PARGi treatment or (**C**) 24–72 h wash-out PARGi treatment, normalised to DMSO control. Mean of ≥3 biological replicates. (**D**) Exemplar sensitive (OVMANA) and resistant (OVSAHO) cell proliferation curves (means of 2 biological replicates). (**E**) AUC from (D) were used to generate proliferative EC_50_ curves shown, mean of 2 biological replicates. (**F**) Proliferative EC_50_s for panel were determined, means of ≥2 biological replicates. PRISM could not accurately calculate EC_50_ for resistant cells, therefore for highly resistant cell lines, EC_50_ was approximated as 50μM (half the maximal concentration tested), and for less highly resistant OV56, EC_50_ was determined manually. (**G**) Colony formation in response to continuous PARPi treatment was quantified, and normalised to DMSO control, mean of ≥3 biological replicates. Statistics: 2-way ANOVA with Dunnett’s multiple comparisons test, selected comparisons were between drug treatments and DMSO control within each cell line. Error bars represent SEM. (**H**) Venn diagram summarising differential sensitivity.

**Figure S2.**
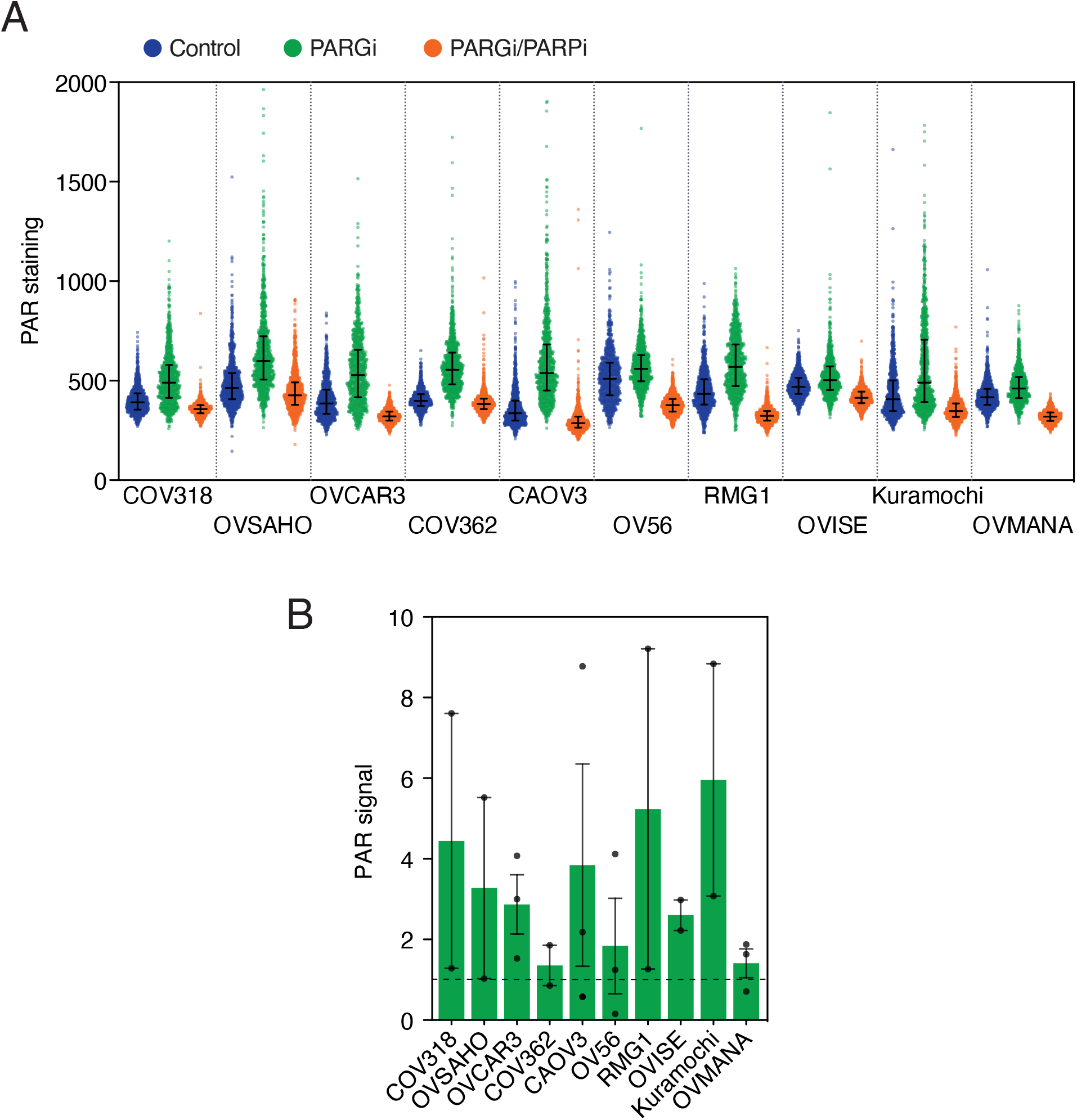
PAR chain levels are comparable in sensitive and resistant cell lines. (**A**) Quantification of PAR staining intensity per cell in 1 biological replicate, 1000 cells shown per condition. (**B**) Graph shows mean quantification of PAR staining by immunoblotting (≥2 biological replicates). Error bars represent SEM. Representative staining images in main figure.

**Figure S3.**
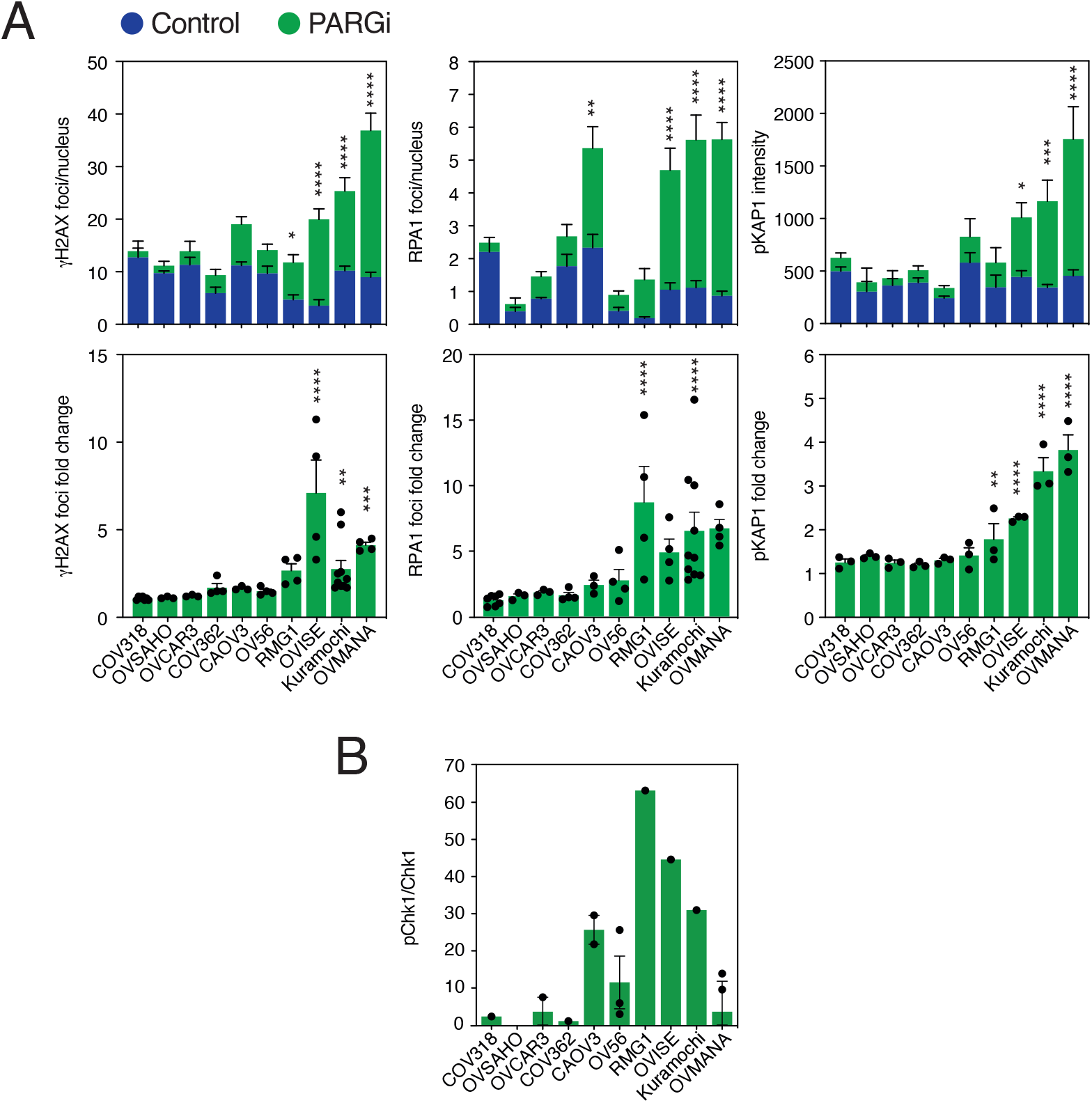
PARGi sensitive cells exhibit markers of replication stress and DNA damage response following PARGi treatment. (**A**) Upper panel: Quantification of foci per nucleus after 48 h PARGi treatment (γH2AX and RPA1), or nuclear pKAP1 intensity after 72 h PARGi treatment; Lower panel: Results from upper panel were normalised to DMSO and represented as a fold-change. Graphs show means of ≥3 biological replicates. (**B**) Quantification of pChk1 immunoblotting shown in main figure, graphs show means of ≥1 biological replicate(s). Data are expressed as the increase resulting from treatment as a percentage of maximum response (achieved with HU) to correct for inter-line variation i.e. % PARGi (pChk1/Chk1)/HU - % DMSO (pChk1/Chk1)/HU. Statistics: 1-way ANOVA with Sidak multiple comparisons test (A), selected comparisons were between PARGi treated values and DMSO control within each cell line. Error bars represent SEM (A, B).

